# Hydrogen Bonding Magnetic Resonance (HBMR)-based Cyclic Electromagnetic DNA Simulation (CEDS) can affect DNA Hybridization and Conformation

**DOI:** 10.1101/2024.08.24.609483

**Authors:** Suk Keun Lee, Dae Gwan Lee, Yeon Sook Kim

## Abstract

The proton magnetic resonance is commonly used in MRI machines with a strong magnetic field of over 1 T, while this study hypothesized that a weak magnetic field below 0.01 T can induce the proton magnetic resonance in hydrogen bonds of double-stranded DNA (dsDNA). This study found the hydrogen bonding magnetic resonance (HBMR) in dsDNA can modulate the conformations and functions of dsDNAs (*1*). Decagonal or dodecagonal cyclic electromagnetic DNA simulation (CEDS) was designed to target three-dimensional structures of randomly oriented dsDNAs with about 25% efficiency. This study found that the most effective magnetic exposure times for inducing an electric potential in A-T and G-C base pairs were 280 and 480 msec, respectively. Decagonal CEDS using a target sequence at 20-25 Gauss for 30 min was able to induce sequence-specific hybridization of target short oligo-dsDNAs in 0.005M NaCl solution and their unique conformation in 0.1M NaCl solution. There was a tendency that Pyu oligo-dsDNAs showed more hybridization and unique conformational changes by CEDS using a target sequence than Puy oligo-dsDNAs. This CEDS effect on Pyu ds6(2C2A) increased with CEDS time up to 90 min and gradually decreased to about half (51.8%) of the increase at 240 min resting time. CEDS influenced the oligo-dsDNAs to increase their infrared (IR) absorbance at approximately 3700-2800 cm^-1^ band compared to the positive and negative controls, which was more dominant in Pyu oligo-dsDNAs than in Puy oligo-dsDNAs. Therefore, it is postulated that HBMR-based CEDS using a weak magnetic field can increase the hybridization potential of oligo-dsDNAs and subsequently lead to unique DNA conformation required for the initiation of various DNA functions. Therefore, it is suggested that decagonal CEDS play a role in regulating the function of target short oligo-dsDNA.

---

Weak magnetic field, 100 Gauss can affect spin-orbital interaction of hydrogen atom and induce minute energy, about 1.2 μeV, which is calculated by Zeeman equation (***E = g***_***l***_***μ***_***B***_***Bm***_***l***_ ; where *g*_*l*_ is orbital gyromagnetic ratio, *μ*_*B*_ is Bohr magneton, *B* is magnetic field strength, and *m*_*l*_ is magnetic quantum number (*m*_*l*_ = 1)(*2-4*)). The electric energy from hydrogen atom is subsequently accumulated in the associated hydrogen bonds, if the quantum radiation were negligible. This study used a weak magnetic field, below 30 Gauss, to affect the hydrogen bonds of oligo-dsDNA and influence the function of oligo-dsDNA.

The cyclic electromagnetic DNA simulation (CEDS) was designed to target hydrogen bonds of dsDNA by rotating a decagonal or dodecagonal magnetic field. CEDS uses five or six pairs of electromagnetic bars aligned in a line facing each other by passing their magnetic field through the center of horizontal magnetic plane. Decagonal and dodecagonal CEDS can target B-type dsDNA and Z-type dsDNA, respectively, depending on the three-dimensional structure of dsDNA (Fig. 1) (*5, 6*).

**Fig. 1.**
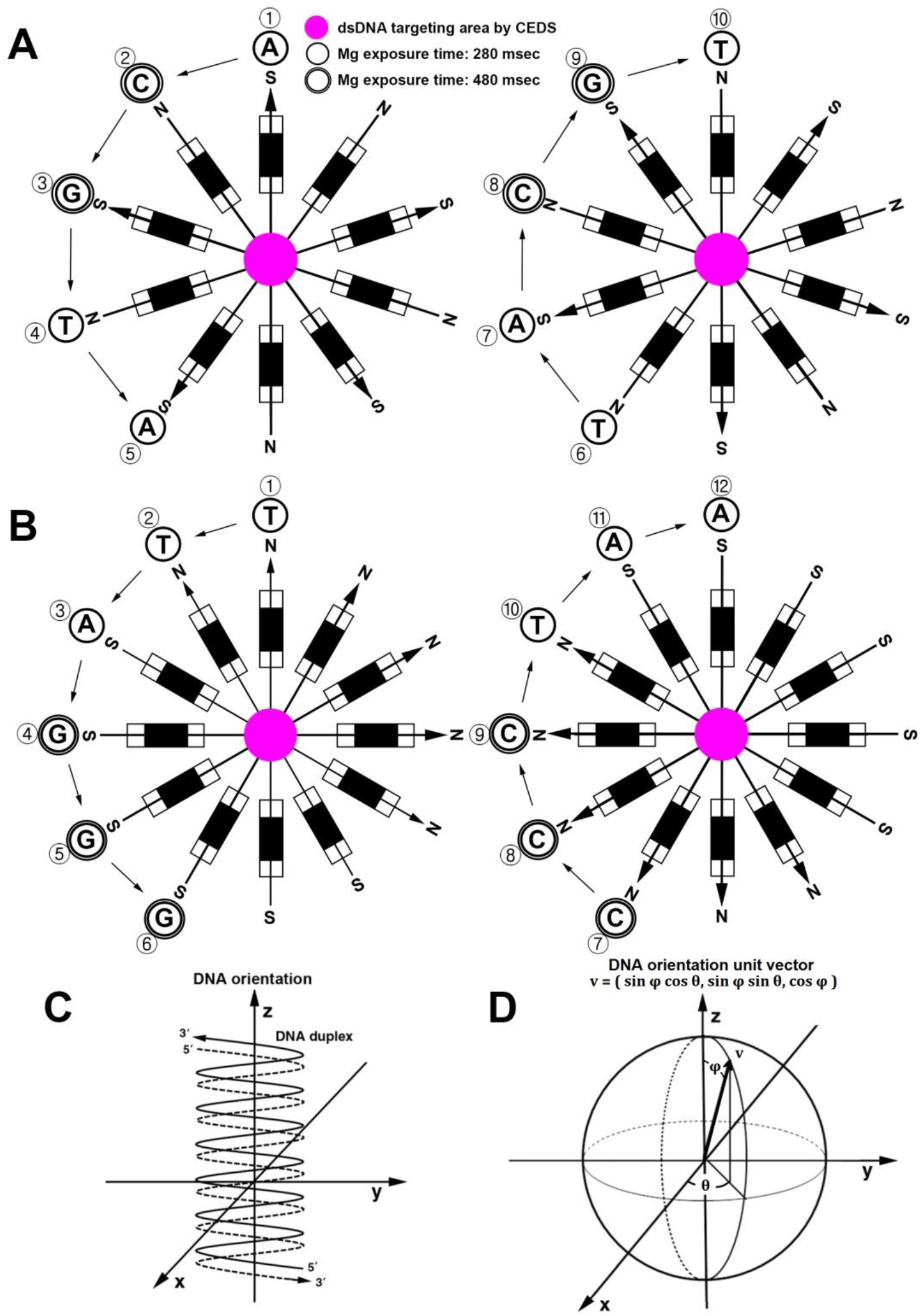
Schematic illustration of CEDS apparatus (A,B). B-type dsACGTA and Z-type dsTTAGGG are targeted by decagonal (A) and dodecagonal (B) CEDS in the procedures of ①→⑩ and ①→ ⑫, respectively. A model of the physical efficiency of CEDS on dsDNAs (C,D). The average physical efficiency of CEDS on randomly distributed dsDNAs in three dimensions is given by 25%.

CEDS uses a magnetic field of 10-30 Gauss and sequentially rotating to simulate the structure of dsDNA for 15-30 min. The direction of magnetic field polarity between the base pairs and for the circular direction of magnetic field are not obligatory due to the chiral structures of dsDNA in three-dimensional space, but should be done sequentially in relation to neighboring base pairs in dsDNA. A-T and G-C base pairs are exposed with different magnetic exposure time, 280 and 480 msec, respectively, according to the determination of effective exposure time for A-T and G-C base pairs (Fig. 2 and 3) (*1*).

**Fig. 2.**
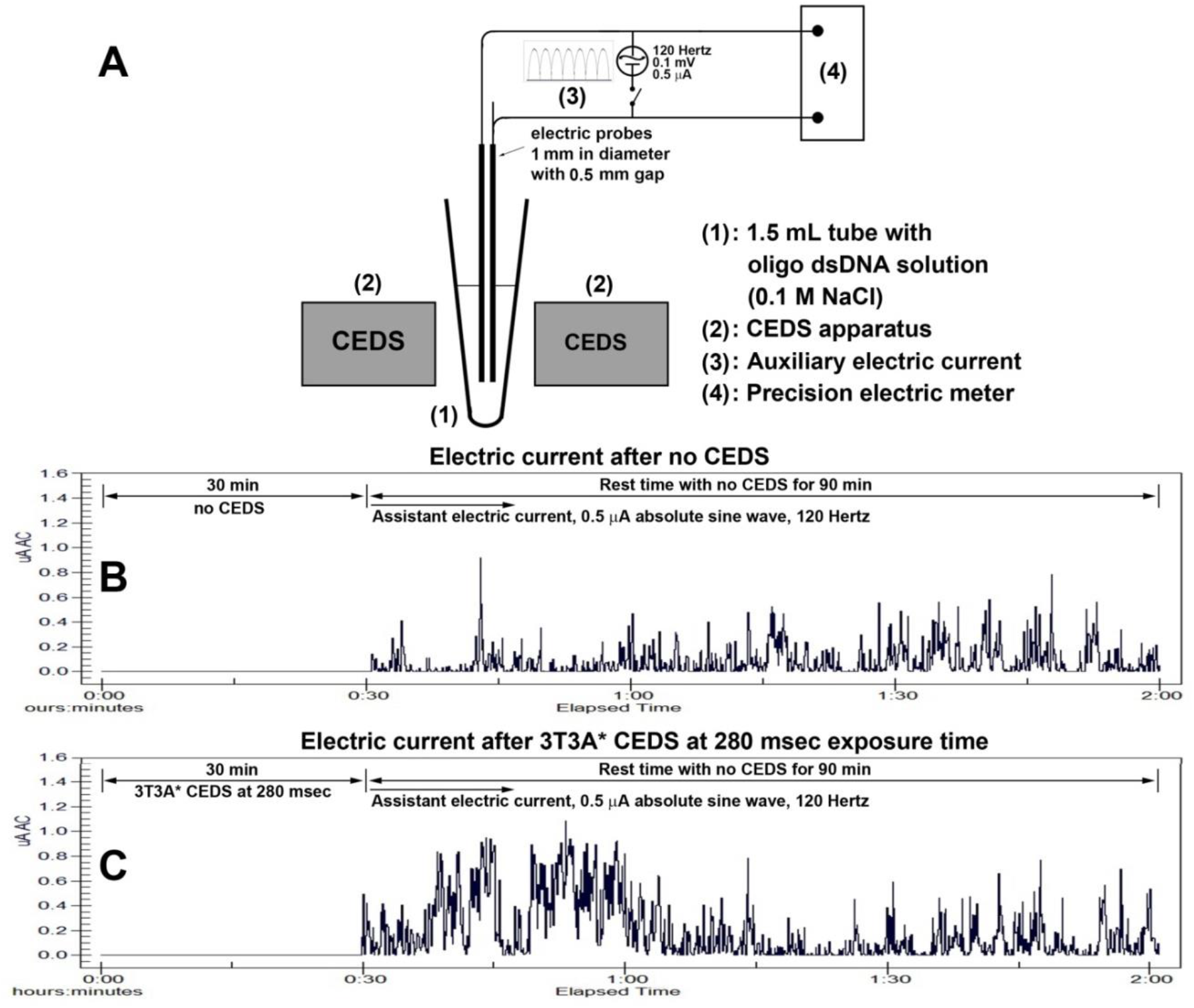
Detection of electric current discharged from oligo-dsDNAs after CEDS treatment. A: Schematic illustration of electric current detection apparatus installed with auxiliary electric current, 0.5 μA absolute sine wave 120 Hertz, between electrodes. It was observed that the electric current increased significantly at 280 msec exposure time for A-T base pair during 40-70 min experimental period (C). On the other hand, the ds3T3A in 0.1 M NaCl solution was not treated with decagonal CEDS for 30 min, there was no increase of electric current during the resting time for 90 min (B).

**Fig. 3.**
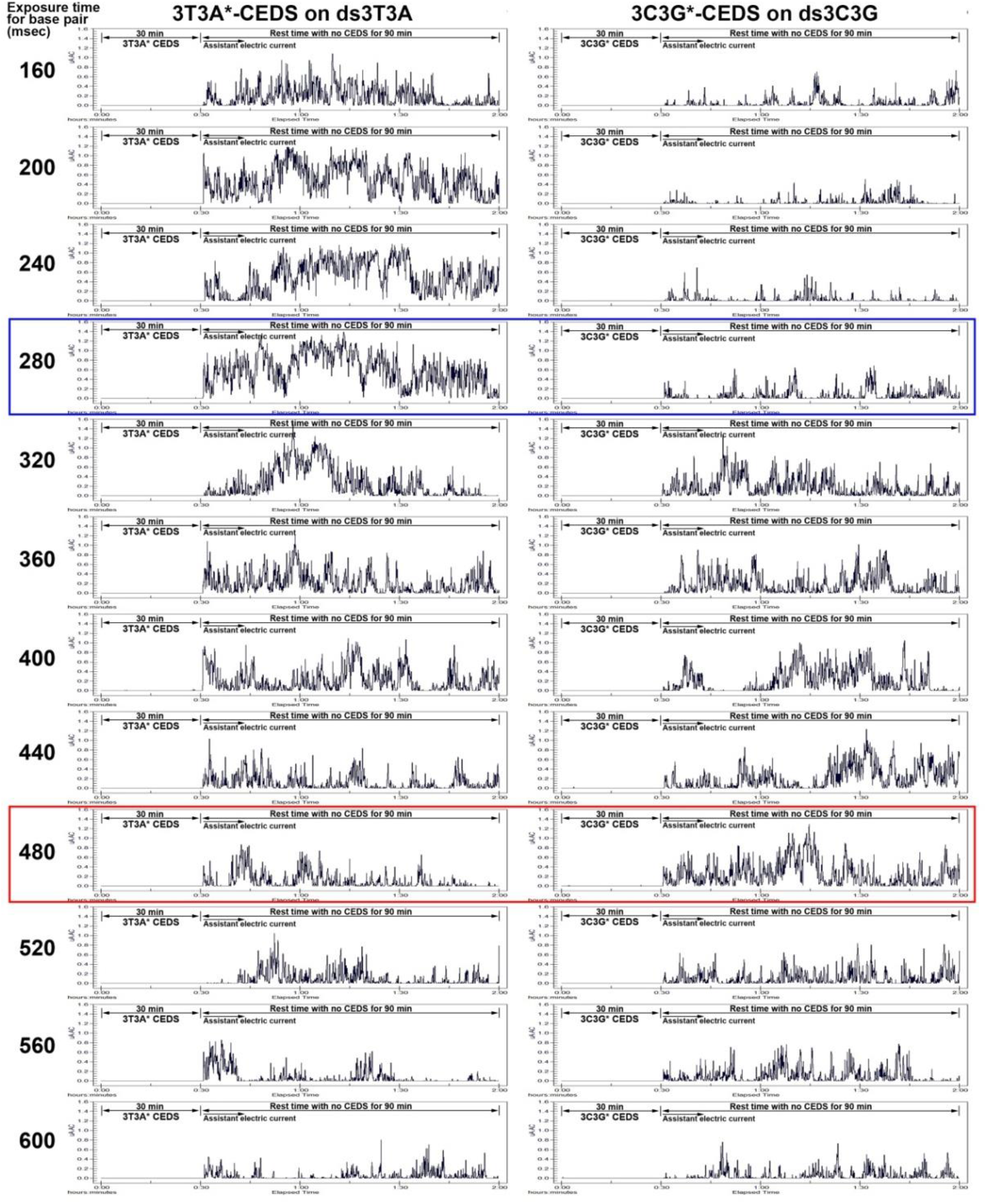
Comparison of electric induction potential in oligo-dsDNAs by decagonal CEDS at different magnetic exposure time, 160-600 msec, between A-T base pairs of ds3T3A and G-C base pairs of ds3C3G. Noted the most dominant increase of electric current in ds3T3A at 280 msec magnetic exposure time contrast in ds3C3G (blue square), and the most dominant increase of electric current in ds3C3G at 480 msec magnetic exposure time contrast in ds3T3A (red square).

## Estimation of the physical efficiency of CEDS on DNA double helix

According to the operating principle of CEDS, the dsDNA in Fig. 1C, which is oriented in the +z axis direction with the sense sequence (solid line) pointing upwards and the antisense sequence (dashed line) pointing downwards, is assumed to be fully affected by the CEDS. However, such exact alignment occurs very rarely because dsDNAs have random orientations in three dimensions. The CEDS is assumed to have no effect on those dsDNAs that are placed horizontally or turned upside down. Taking these cases into consideration, the physical efficiency of CEDS on a dsDNA oriented in the (θ,φ) direction can be modeled as

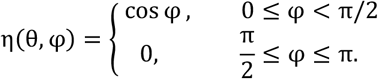

In particular, the case φ = 0 corresponds to the dsDNAs that are oriented in the +z axis direction, and therefore are affected by the CEDS with full efficiency, η(θ, φ)=1. The cases φ = π/2 and φ = πcorrespond to the dsDNAs that are placed horizontally or turned upside down, respectively, and therefore are affected by the CEDS with efficiency η(θ, φ)=0, indicating no effect by the CEDS.

By denoting the orientation of a dsDNA as a unit vector in the spherical coordinate system (0≤θ<2π, 0≤φ≤π, r=1), the random orientations of dsDNAs in three dimensions is represented by the uniform distribution over the unit sphere. Therefore, the average physical efficiency of CEDS on randomly distributed dsDNAs in three dimensions is given by

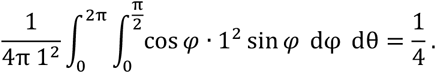

In summary, the theoretical efficiency of CEDS on randomly distributed dsDNAs in three dimensions is limited by 25% (Fig. 1 C,D).

## Determination of magnetic exposure time for A-T and G-C pairs

According to the HBMR hypothesis, the repeated CEDS can accumulate electrostatic charge in the bases of target dsDNA, which is subsequently discharged into buffer solution slowly. To accelerate the electrical discharge, an auxiliary electrical current was installed between the electrical probes, and then we were able to measure the weak electrical discharge from oligo-dsDNAs. To determine the optimal magnetic exposure time on A-T and G-C base pairs, a series of experiments was performed (Fig. 2).

Two electrical probes, each approximately 1 mm in diameter, were placed 0.5 mm apart in a 1.5 mL tube (Fig. 2 A). The ds3T3A (100 pmole/μL) in 0.1 M NaCl solution was treated with decagonal 3T3A*-CEDS using a 13-15 Gauss magnetic field for 30 min. The electric current was then measured under the stimulation of an auxiliary electric current by applying 0.5 μA absolute sine wave at 120 Hz during the resting period for 90 min. It was observed that the electric current increased significantly at 280 msec exposure time for A-T base pair during the 40-70 minute experimental period (Fig. 2 C). On the other hand, the ds3T3A in 0.1 M NaCl solution was not treated with CEDS for 30 min, there was no significant increase of electric current during the resting time for 90 min (Fig. 2 B). These data may indicate that CEDS effect on dsDNA could be evaluated by measuring the electric current discharged from dsDNA after CEDS treatment.

In order to determine the optimal magnetic exposure time on the A-T and G-C base pairs of dsDNA, a series of experiments were performed using the same methods as above to compare the amount of electric current discharge from ds3T3A and ds3C3G after decagonal 3T3A*-CEDS and 3G3C*-CEDS, respectively, at different magnetic exposure times, 160, 200, 240, 280, 320, 360, 400, 440, 480, 520, 560, and 600 msec. As a result, ds3T3A, which is composed of only A-T base pairs, showed the dominant increase of electric current discharge at 240 -280 msec magnetic exposure time compared to ds3C3G, which is composed of only G-C base pairs. On the other hand, ds3C3G showed the dominant increase of the releasing electric current at 440-480 msec magnetic exposure time compared to ds3T3A (Fig. 3).

As a result, ds3T3A consisting of A-T base pairs showed dominant electric discharge at 240 -280 msec of magnetic exposure time after decagonal CEDS compared to ds3C3G consisting of G-C base pairs, while ds3C3G showed dominant electric discharge at 440-480 msec compared to ds3T3A (Fig. 3). The data suggest that A-T base pair polarity is more saturated by CEDS at 240-280 msec of magnetic exposure time than G-C base pair polarity, and G-C base pair polarity is more saturated by CEDS at 440-480 msec than A-T base pair polarity. Therefore, the present CEDS used the magnetic exposure times of 280 msec and 480 msec for A-T and G-C base pairs, respectively.

## CEDS effect on the hybridization of oligo-dsDNAs

### 1) Equilibrium state between oligo-dsDNA and oligo-ssDNA pair in 0.005M NaCl solution

Pyu ss2(3C3A) and ss2(3T3G), were purchased and dissolved in 0.2M, 0.05M, 0.01M, and 0.005M NaCl solution to obtain 200 μL mixture of ss2(3C3A) and ss2(3T3G) at 100 pmole/μL. After 10 min at room temperature, each sample was analyzed by HPLC using a reverse-phase silica column with different concentrations of mobile phase, 0.2M, 0.05M, 0.01M, and 0.005M NaCl, at a flow rate of 0.3 mL/min, UV260. At room temperature, a pair of ss2(3C3A) and ss2(3T3G) spontaneously hybridized in 0.2M and 0.05M NaCl solution, respectively, producing dominant dsDNA peaks, while in 0.005M NaCl solution, an equilibrium between dsDNAs and ssDNAs appeared (Fig. 4A).

**Fig. 4.**
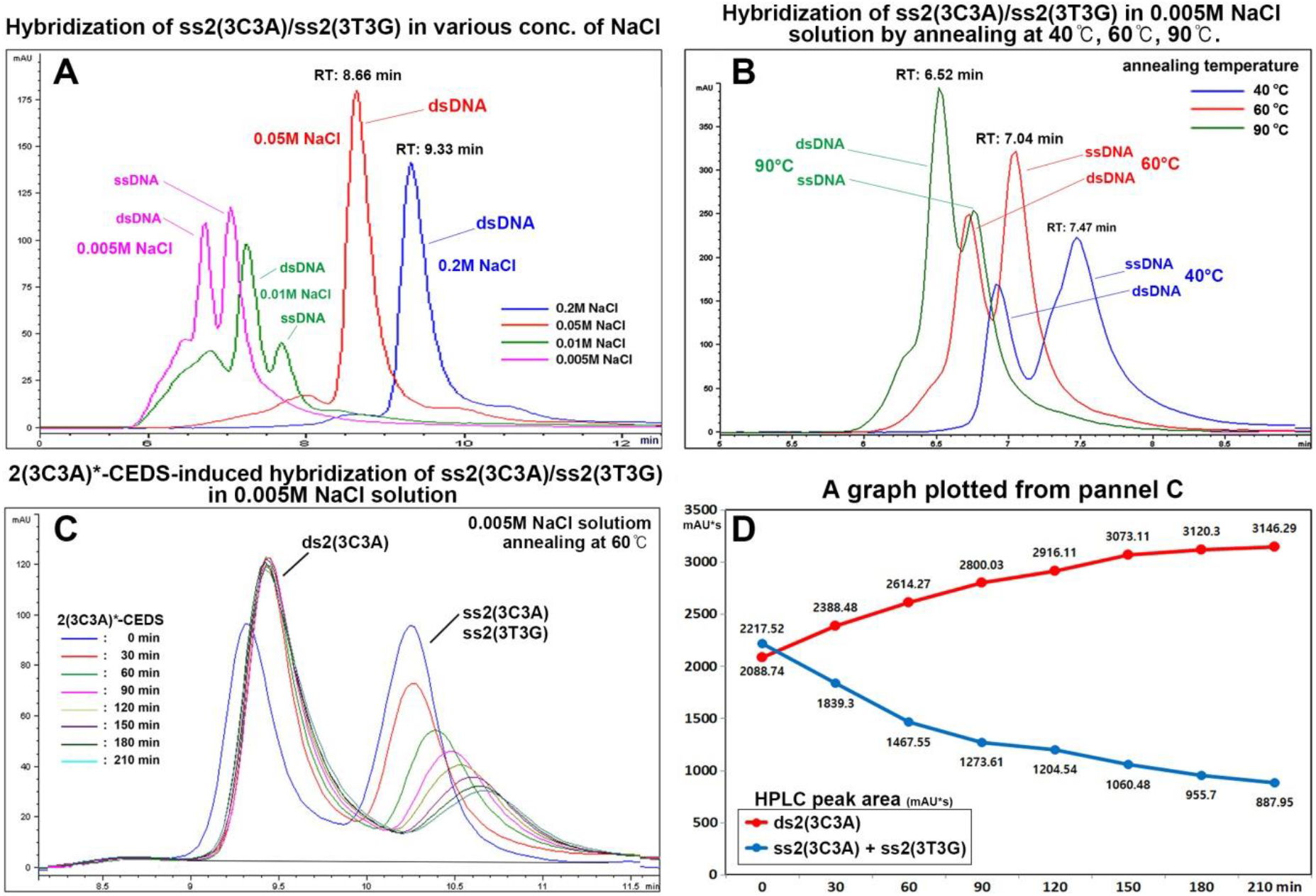
HPLC analysis of oligo-Pyu dsDNA hybridization status depending on NaCl concentration. A pair of oligo-ssDNAs, ss2(3C3A) and ss2(3T3G), was strongly hybridized in 0.05M and 0.2M NaCl solutions at room temperature, while loosely hybridized in 0.005M and 0.01M NaCl solutions (A). This state of equilibrium was most prominent when annealed at 60°C compared to 40°C and 90°C (B). When a pair of ss2(3C3A) and ss2(3T3G) was dissolved in a 0.005M NaCl solution and annealed at 60°C, the ssDNA pairing was then hybridized by decagonal 2(3C3A)*-CEDS depending on CEDS time, 30, 60, 90, 120, 150, 180, and 210 min (C). A graph was created using panel C data (D). RT: Retention time.

CEDS resulted in a shift in the balance between double-stranded DNAs (dsDNAs) and single-stranded DNAs (ssDNAs), with a prominent peak area of dsDNAs and a small peak area of ssDNAs. The shift in the balance between double-stranded DNAs (dsDNAs) and single-stranded DNAs (ssDNAs) was dependent on CEDS time, as shown in Fig. 4C. A plotted graph of the data clearly shows a gradual increase in the dsDNA peak for ds2(3C3A), and a gradual decrease in the ssDNA peak for ss2(3C3A) and ss(3T3G), also dependent on CEDS time (Fig. 4D). The data indicate that 2(3C3A)*-CEDS induced hybridization of a oligo-ssDNA pair, ss2(3C3A) and ss2(3T3G), dissolved in 0.005M NaCl solution, resulted in a dominant increase of ds2(3C3A) peak depending on CEDS time compared to ss2(3C3A)/ss2(3T3G) peak.

Meanwhile, ss2(3C3A) and ss2(3T3G) were dissolved in 0.005 M NaCl solutions and heated to 40°C, 60°C, or 90°C, then gradually cooled to room temperature, and followed by HPLC as described above. The results showed a relatively even equilibrium between dsDNAs and ssDNAs with less tertiary structure at 60°C compared to 40°C and 90°C (Fig. 4B). Therefore, we conducted the hybridization of oligo-dsDNA by dissolving in 0.005M NaCl solution and annealing at 60°C.

A pair of ss2(3C3A) and ss2(3T3G) in 0.005M NaCl was annealed at 60°C as previously described, and treated with decagonal 2(3C3A)*-CEDS at 20-25 Gauss for 210 min. The sample was analyzed by HPLC using the method described above. 2(3C3A)*-CEDS gradually increased the peak area of ds2(3C3A) and decreased the peak area of ss2(3C3A)/ss2(3T3G) depending on CEDS time, indicating that 2(3C3A)*-CEDS induced hybridization of oligo-ssDNA pair in 0.005M NaCl solution, resulted in a dominant increase of ds2(3C3A) peak depending on CEDS time compared to ss2(3C3A)/ss2(3T3G) peak (Fig. 4D).

### 2) Pyu oligo-dsDNAs in 0.005M NaCl solution were hybridized by CEDS and reverted to the normal state in 18 hours

The effect of CEDS on hybridization potential of oligo-dsDNAs could be clearly observed in low salt solution of 0.005M NaCl. Pyu ds2(6C6A), ds3(4C4A), ds4(3C3A), ds6(2C2A), ds12(CA), and ds12C12A in 0.005M NaCl solution were prepared using the method described above. The sample was then treated with decagonal CEDS using a target sequence at 20-25 Gauss for 30 min, at room temperature and analyzed by HPLC as previously described.

As a result, ds3(4C4A), ds4(3C3A), ds6(2C2A), and ds12(CA) hybridized more strongly by CEDS than untreated controls (Fig. 5 C,E,G,I), and these DNA hybridizations reverted to the normal state of untreated controls in 18 h (Fig. 5 D,F,H,J). On the other hand, ds2(6C6A) and ds12C12A appeared to hybridize spontaneously but incompletely in 0.005 M NaCl solution, probably due to the formation of multiple G-quadruplex complexes. However, the samples hybridized more by CEDS than untreated controls (Fig. 5 A,B,K,L). The data show that ds3(4C4A), ds4(3C3A), ds6(2C2A), and ds12(CA) in 0.005M NaCl solution hybridized more by CEDS than untreated controls, but eventually returned to the normal state within 18 hours.

**Fig. 5.**
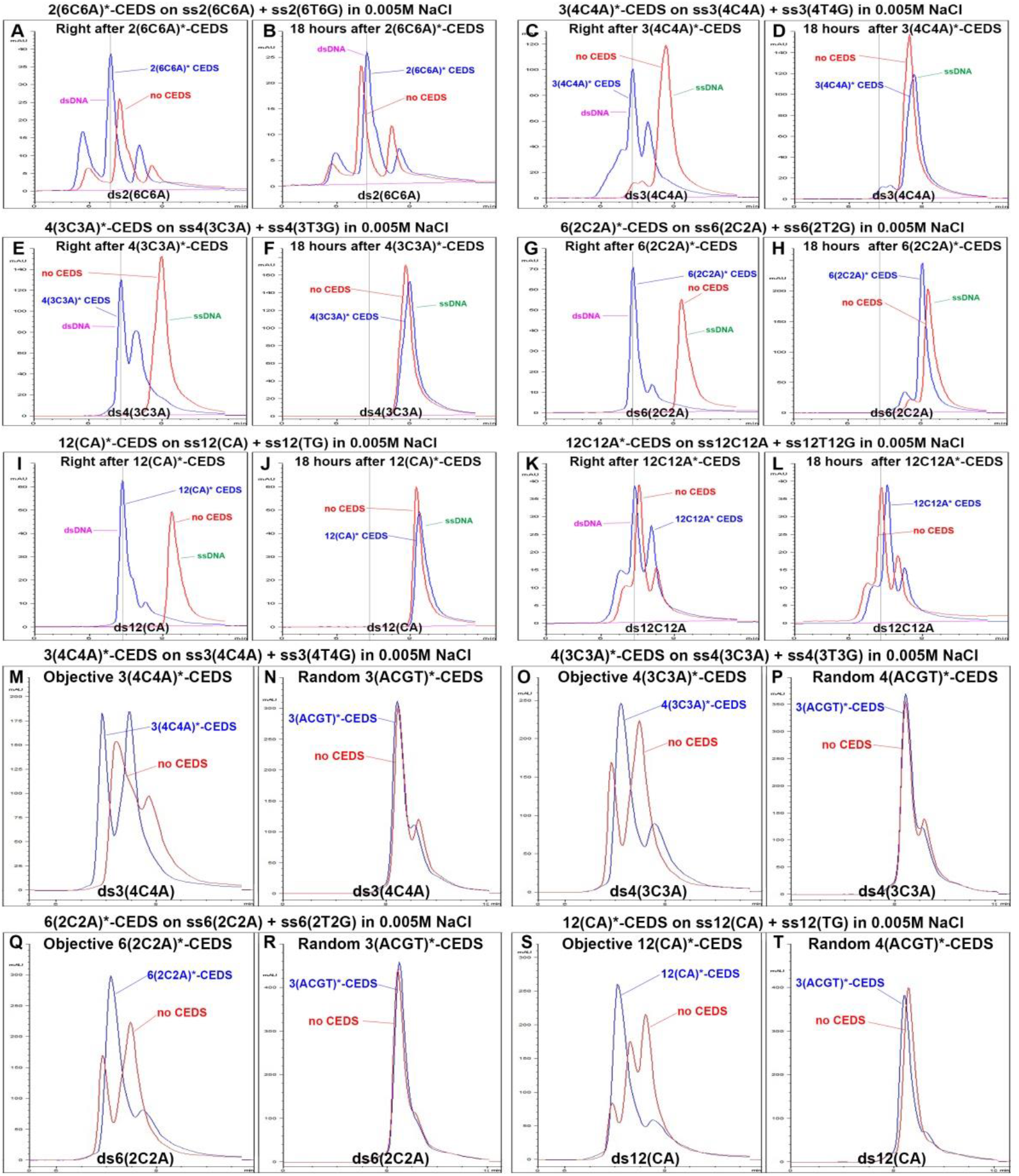
CEDS-induced hybridization of ds2(6C6A) (A,B), ds3(4C4A) (C,D,M,N), ds4(3C3A) (E,F,O,P), ds6(2C2A) (G,H,Q,P), ds12(CA) (I,J,S,T), ds12C12A (K,L) in 0.005M NaCl solution with 60°C annealing.

### 3) Pyu oligo-dsDNAs in 0.005M NaCl solution were hybridized by CEDS in a sequence-specific manner

Pyu ds3(4C4A), ds4(3C3A), ds6(2C2A), ds12(CA) in 0. 005M NaCl solution were prepared using the method described above and assessed their hybridization by decagonal CEDS using each sequence. The CEDS resulted dominant peaks of ds3(4C4A), ds4(3C3A), ds6(2C2A), or ds12(CA) compared to untreated controls (Fig. 5 M,O,Q,S), whereas the random sequence*-CEDS (3(ACGT)*-CEDS) induced incomplete hybridization peaks (Fig. 5 N,P,R,T). The data suggest that oligo-dsDNAs were hybridized by the target sequence*-CEDS more than by 3(ACGT)*-CEDS.

Taken together, it is evident that the target sequence*-CEDS facilitates the hybridization of oligo-dsDNAs in 0.005M NaCl solution depending on CEDS time up to 210 min and in a sequence-specific manner when compared to 3(ACGT)*-CEDS. In addition, the CEDS-induced hybridization is reversible to the normal state within 18 h.

### 4) The conformational changes of ds12/n(nCnA) (n=1,2,3,4,6,12) in 0.1M NaCl solution by 12/n(nCnA)*-CEDS or 12A*-CEDS

A pair of ss12/n(nCnA) and ss12/n(nTnG) (n=1,2,3,4,6,12) was purchased and dissolved in 0.1M NaCl solution at 100 pmole/μL by heating to 60°C and slowly cooling to room temperature, and then ds12/n(nCnA) hybridized without denaturation caused by overheating. The oligo-dsDNA samples were treated with dodecagonal CEDS using each DNA sequence at 20-25 Gauss for 30 min at room temperature, and immediately followed by HPLC using a reverse phase column packed with silica beads and 0.1M NaCl running buffer at 30°C (Fig. 6).

**Fig. 6.**
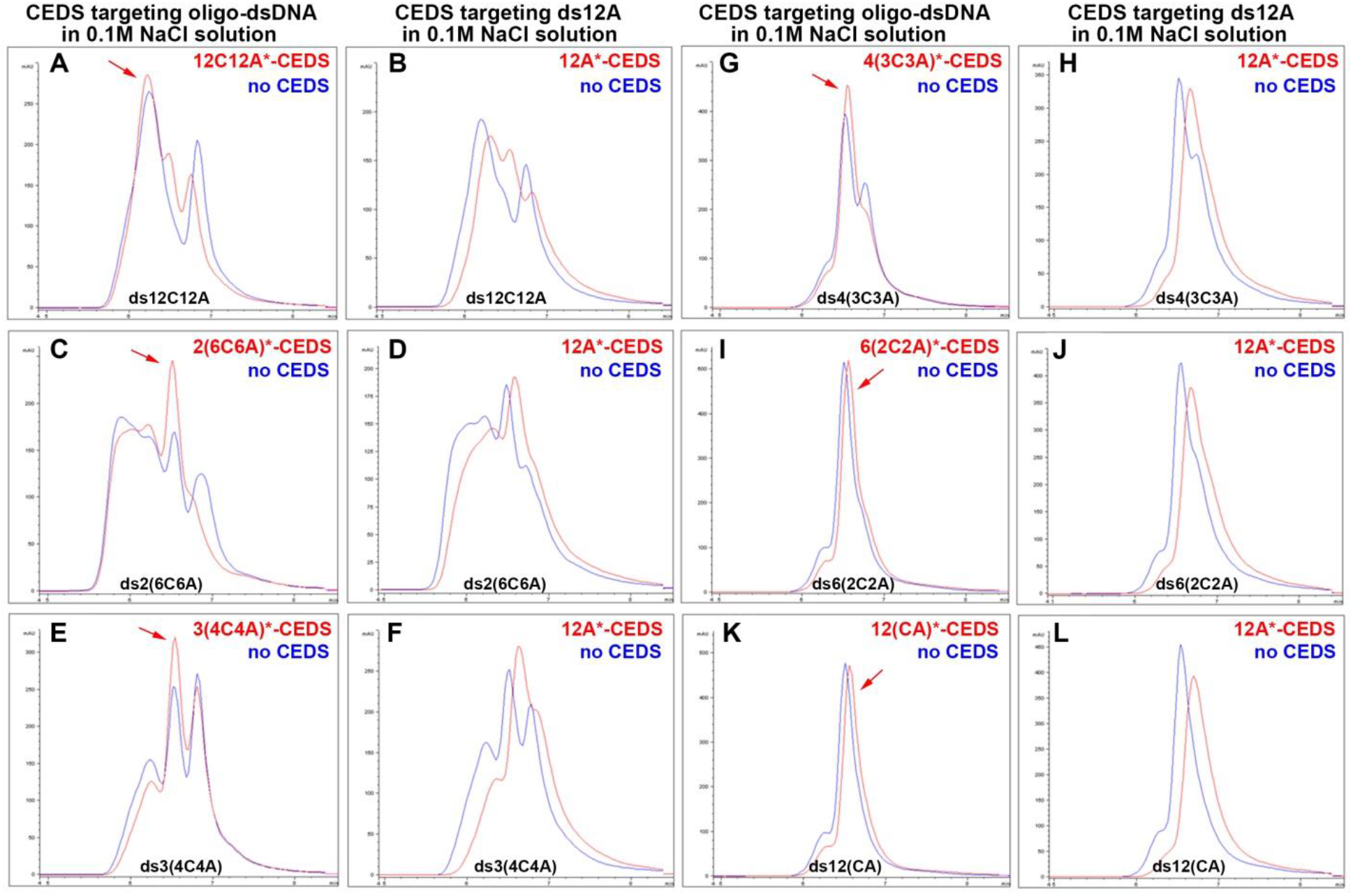
HPLC analysis for the conformational changes of ds12/n(nCnA) (n=1,2,3,4,6,12) in 0.1M NaCl solution by dodecagonal 12/n(nCnA)*-CEDS or 12A*-CEDS. A dominant peak of oligo-dsDNA (red arrow).

In the HPLC results, ds12/n(nCnA) (n=1,2,3,4,6,12) in 0.1M NaCl solution showed different patterns of peak area depending on the number of DNA segments. The ds6(2C2A) and ds12(CA), which have 6 and 12 segments, respectively, showed a single dominant peak, while ds12C12A, ds2(6C6A), ds3(4C4A), and ds4(3C3A), which have 1, 2, 3, and 4 segments, respectively, showed multiple peaks. In particular, ds2(6C6A) and ds12A12C showed an anomalous peak with an early retention time, indicating strong aggregation of DNA, probably G-quaruplex (Fig. 6 A-D).

Dodecagonal CEDS targeting ds12/n(nCnA) (n=1,2,3,4,6,12) usually increased the dominant peak of oligo-dsDNA and decreased its retention time compared to untreated controls. This CEDS effect was strong in ds3(4C4A) and ds4(3C3A), weak in ds12C12A and ds2(6C6A), and almost absent in ds6(2C2A) and ds12(CA), which tended to show delayed retention times. On the other hand, 12A*-CEDS was simultaneously performed as a positive control, and it was found that 12A*-CEDS tended to deform or reduce the dominant peaks of oligo-dsDNAs and increase their retention time, contrary to 12/n(nCnA)*-CEDS (Fig. 6).

## CEDS effect on the DNA conformation of oligo-dsDNAs in 0.1M NaCl solution

### 1) Simple Pyu oligo-dsDNAs showed larger HPLC peak by CEDS than simple Puy oligo-dsDNAs

Simple Pyu and Puy oligo-dsDNA consisting of A-T or G-C pairs only, including ds3T3A, ds3A3T, ds3C3G, ds3G3C, ds3C3A, ds3A3C, were dissolved in 0.1M NaCl solution and annealed at 90°C and slowly cooled at room temperature. The samples were treated with decagonal CEDS using each sequence for 30 min and analyzed by HPLC using the method described above.

Pyu ds3T3A showed 4.8% increase in peak area by 3T3A*-CEDS, and Puy ds3A3T showed 0.8% decrease by 3A3T*-CEDS, while 12A*-CEDS resulted in 6.9% and 2.2% increase, respectively (Fig. 7 A,B). Pyu ds3C3G showed 5.2% increase in peak area by 3C3G*-CEDS, and Puy ds3G3C showed 10.4% increase by 3G3C*-CEDS, while 12A*-CEDS resulted in 12.1% and 13.1% increase, respectively (Fig. 7 C,D). Pyu ds3C3A showed 0.7% decrease in peak area by 3C3A*-CEDS, and Puy ds3A3C showed 2.2% decrease by 3A3C*-CEDS, while 12A*-CEDS resulted in 0.9% and 7.5% decrease, respectively (Fig. 7 E,F). The data indicate that Pyu oligo-dsDNAs tend to be more responsive to CEDS than Puy oligo-dsDNAs.

**Fig. 7.**
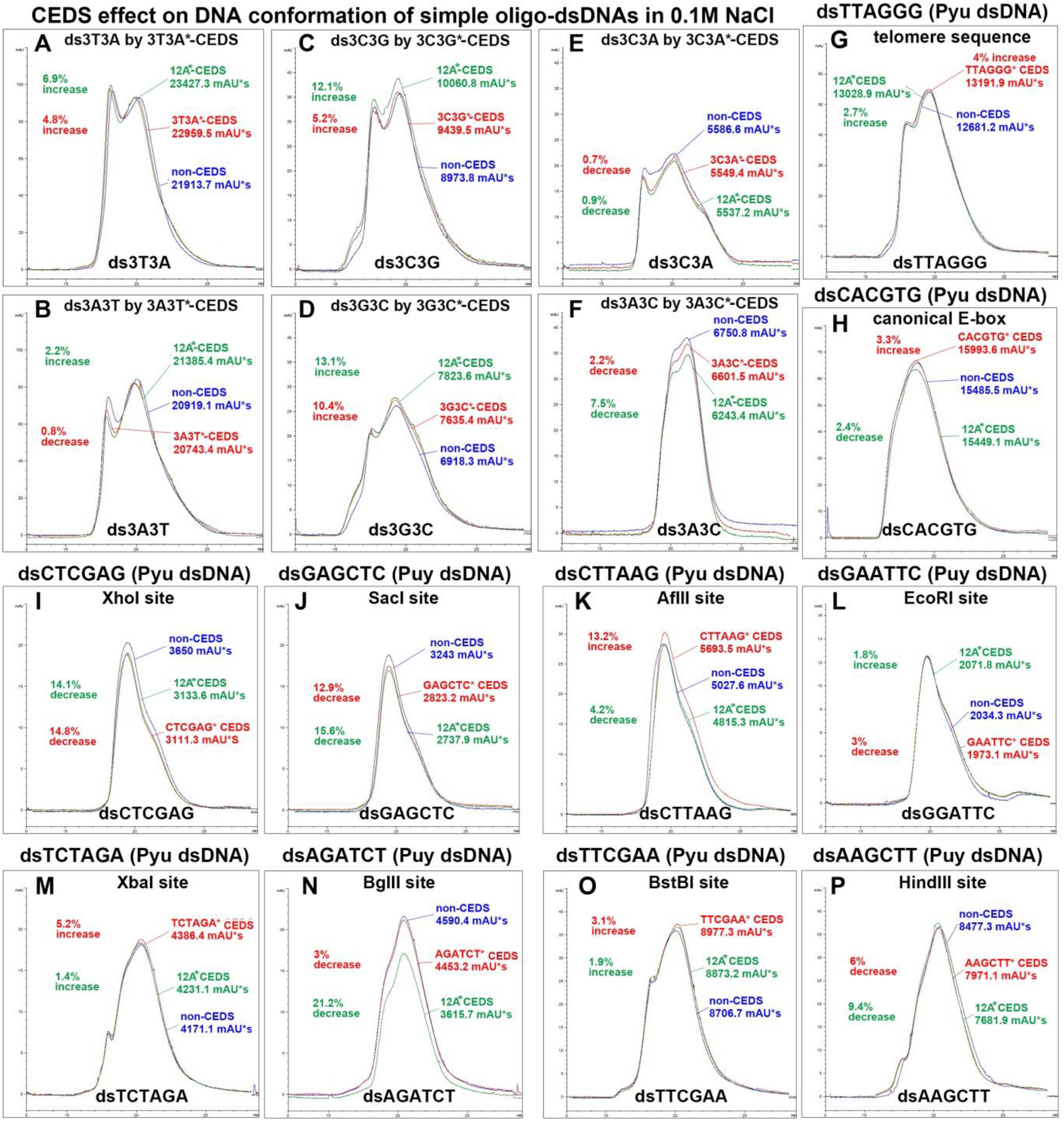
HPLC for CEDS-induced conformational changes between simple Pyu/Puy dsDNAs, including ds3T3A/ds3A3T (A,B), ds3C3G/ds3G3C (C,D), and complex Pyu/Puy dsDNAs, including ds3C3A/ds3A3C (E,F), dsTTAGGG/dsCACGTG (G,H), dsCTCGAG/dsGAGCTC (I,J), dsCTTAAG/dsGAATTC (K,L), dsTCTAGA/dsAGATCT (M,N), and dsTTCGAA/dsAAGCTT (O,P) in 0.1M NaCl solution.

### 2) Complex Pyu oligo-dsDNAs tended to have larger HPLC peak by CEDS than complex Puy oligo-dsDNAs

Six bps complex oligo-dsDNAs of telomere repeat sequence, canonical E-box sequence, and restriction endonuclease (RE) binding site sequences including XhoI, SacI, AfIII, EcoRI, XbaI, BgIII, BstBI, and HindIII, consisting of both A-T and G-C pairs, were prepared in 0.1M NaCl solution at 100 pmole/μL using the method described above. The samples were treated with decagonal CEDS using each sequence at 20-25 Gauss for 30 min and analyzed by HPLC as previously described.

The telomere repeat sequence, Pyu dsTTAGGG, showed 4% increase in peak area by TTAGGG*-CEDS, and the canonical E-box sequence consisting of three Pyu dsDNA segments, Pyu dsCACGTG, showed 3.3% increase by CACGTG*-CEDS, while 12A*-CEDS resulted in 2.7% increase and 2.4% decrease, respectively (Fig. 7 G,H).

The palindromic Pyu dsCTCGAG (XhoI), dsCTTAAG (AfIII), dsTCTAGA (Xba1), and dsTTCGAA (BstBI) showed 14.8% decrease, 13.2% increase, 5.2% increase, and 3.1% increase in peak area by CTCGAG*-CEDS, CTTAAG*-CEDS, TCTAGA*-CEDS, and TTCGAA*-CEDS, respectively, while 12A*-CEDS resulted in 14.1% decrease, 4.2% decrease, 1.4% increase, and 1.9% increase, respectively (Fig. 7 I,K,M,O).

On the other hand, the palindromic Puy dsGAGCTC (SacI), dsGGATTC (EcoRI), dsAGATCT (BgIII), and dsAAGCTT (HindIII) showed 12. 9%, 3%, 3%, and 6% decrease in peak area by GAGCTC*-CEDS, GGATTC*-CEDS, AGATCT*-CEDS, and AAGCTT*-CEDS, respectively, while 12A*-CEDS resulted in 15.6% decrease, 1.8% increase, 21.2% decrease, and 9.4% decrease, respectively (Fig. 7 J,L,N,P).

In the same line, 12/n(nCnA)*-CEDS can influence ds12/n(nCnA) (n=1,2,3,4,6,12) in 0.1M NaCl solution by increasing the dominant peak of oligo-dsDNA and decreasing its retention time compared to untreated controls (Supplementary Text, Fig. 6). Taken together, it was found that CEDS significantly altered the conformation of target oligo-dsDNAs in a sequence-specific manner. The simple and complex Pyu dsDNAs in 0.1M NaCl solution showed a tendency to increase their conformation by CEDS more than Puy dsDNAs, and these changes were much in contrast to 12A*-CEDS.

### 3) CEDS-treated Pyu oligo-dsDNAs showed gradual increase of HPLC peak area depending on CEDS time in a sequence-specific manner

The CEDS-induced sequential changes of peak area of oligo-dsDNAs were assessed by HPLC analysis. Simple ds6(2T2A) in 0.1M NaCl solution was prepared using the method described above, and treated with the target sequence*-CEDS (6(2T2A)*-CEDS), random sequence*-CEDS (6(ACGT)*-CEDS), or nonspecific sequence*-CEDS (24A*-CEDS) in a dodecagonal fashion at 20-25 Gauss, room temperature for 120 min. During the CEDS, 50 μL of sample was obtained, and analyzed by HPLC as previously described. 6(2T2A)*-CEDS increased the peak area of ds6(2T2A) depending on CEDS time, up to 120 min. Conversely, 6(ACGT)*-CEDS exhibited only a weak increase in the peak area of ds6(2T2A) during CEDS time, while 24A*-CEDS exhibited a slight decrease (Fig. 8A).

**Fig. 8.**
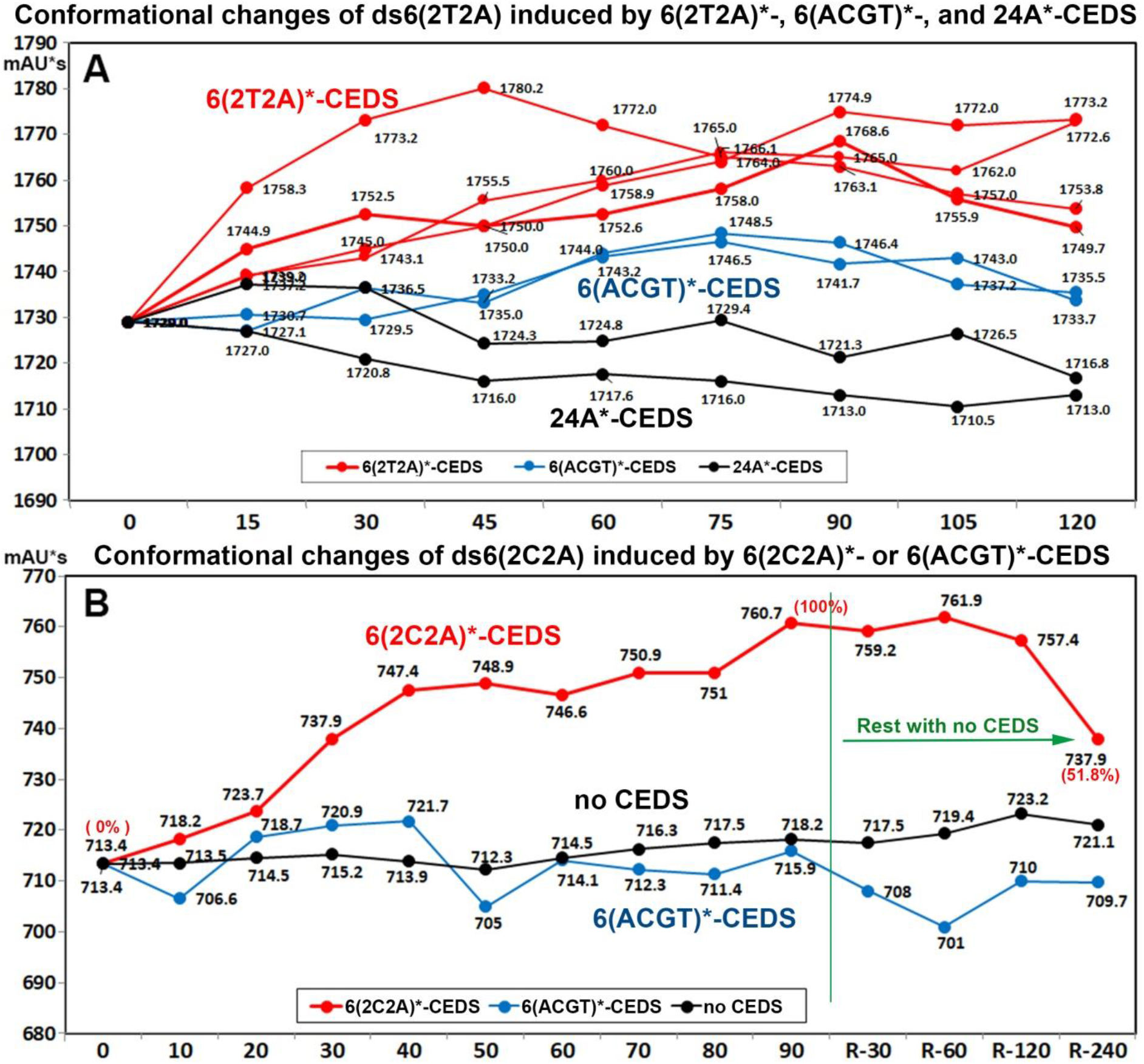
HPLC analysis for the sequential conformational changes of simple ds6(2T2A) and complex ds6(2C2A) induced by ds6(2T2A)*-CEDS and ds6(2C2A)*-CEDS for 120 min and 90 min, respectively, compared to those induced by 6(ACGT)*-CEDS or 24A*-CEDS.

On the other hand, complex ds6(2C2A) in 0.1M NaCl solution was prepared and treated with 6(2C2A)*-CEDS or 6(ACGT)*-CEDS in a dodecagonal fashion for 90 min using the method described above, and the samples were allowed to rest for 240 min. 50 μL of sample was obtained during the CEDS for 90 min, and after the CEDS for 240 min. Each sample was analyzed by HPLC using the method described above.

During the CEDS, 6(2C2A)*-CEDS consistently increased the peak area of ds6(2C2A) until 90 min, and it showed a plateau during 120 min of resting time with minimal change. Thereafter, the peak area showed a gradual decrease, reaching approximately half of its maximum increase (51.8%) at 240 min. In contrast, 6(ACGT)*-CEDS produced irregular increases or decreases in the peak area during the CEDS and resting time, while the untreated control showed minimal change (Fig. 8B).

Both ds3(4C4A) and ds4(3C3A) in 0.2M NaCl solution also showed gradual increase of peak area by 3(4C4A)*-CEDS and 4(3C3A)*-CEDS during the CEDS time of 90 min, while ds3(4C4A) was almost not changed by 12A*-CEDS (Supplementary Text, Fig. 9). The results indicate CEDS can target and influence oligo-dsDNAs dissolved not only in the lower salt solution, 0.1M NaCl, but also in the higher salt solution, 0.2M NaCl, than the normal saline, 0.154M NaCl, in a sequence-specific manner.

**Fig. 9.**
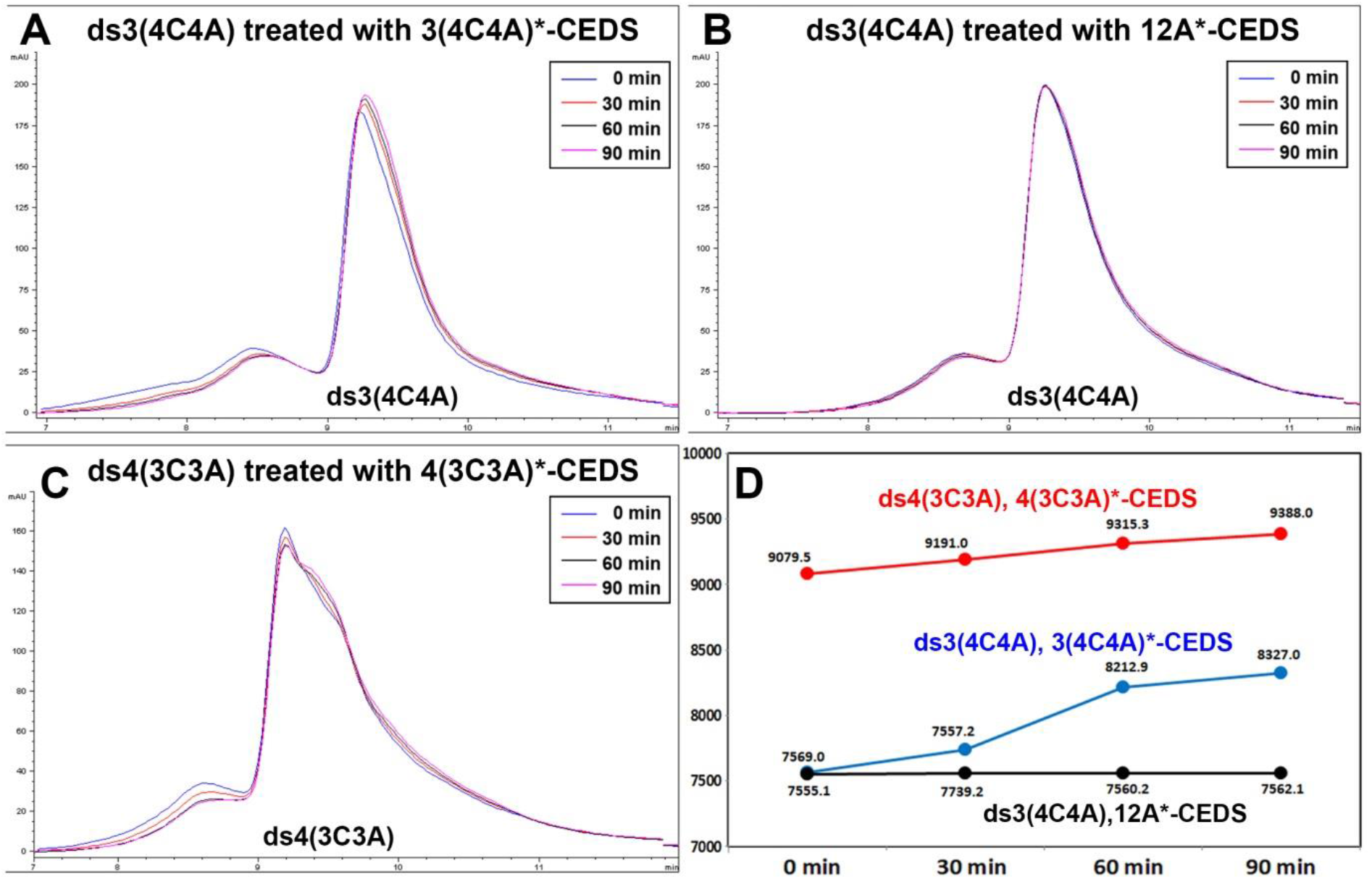
HPLC analysis for the conformational changes of ds3(4C4A) and ds4(3C3A) in 0.2M NaCl solution by dodecagonal 3(4C4A)*-CEDS (A) and 4(3C3A)*-CEDS (C), respectively, for 90 min. B: ds3(4C4A) treated with 12/a-CEDS as a positive control. D: A graph plotted from the data of pannel A-C.

### 4) The conformational changes of ds3(4C4A) and ds4(3C3A) in 0.2M NaCl solution by 3(4C4A)*-CEDS and 4(3C3A)*-CEDS, respectively

The pairs of ss3(4C4A) and ss3(4T4G), ss4(3C3A) and ss4(3T3G) were separately dissolved in 0.2M NaCl solution and annealed at 60°C for 5 min and slowly cooled to room temperature. The ds3(4C4A) and ds4(3C3A) samples were treated with dodecagonal 3(4C4A)*-CEDS and 4(3C3A)*-CEDS, respectively, at 20-25 Gauss for 30 min at room temperature and immediately followed by HPLC using a reverse phase column packed with silica beads and 0.2M NaCl solution running buffer at 30°C. The ds3(4C4A) sample was also treated with dodecagonal 12A*-CEDS as a positive control and simultaneously analyzed by HPLC (Fig. 9).

Since normal saline has 0.154M NaCl concentration, it may be implicative to know the CEDS effect on oligo-dsDNA in 0.2M NaCl solution, which is higher concentration than normal saline. Resultantly, HPLC results showed both ds3(4C4A) and ds4(3C3A) in 0.2M NaCl solution showed gradual increase of peak area by dodecagonal 3(4C4A)*-CEDS and 4(3C3A)*-CEDS during the CEDS time of 90 min, while ds3(4C4A) was almost not changed by 12A*-CEDS (Fig. 9). The results indicate the CEDS can target and influence oligo-dsDNAs dissolved in the higher salt solution, 0.2M NaCl, than the normal saline in a sequence-specific manner.

## Oligo-dsDNAs show the increase in IR absorbance at 3700-2800 cm^-1^ band by CEDS

The N-H stretching of hydrogen bonds of oligo-dsDNA was detected by FT-IR spectroscopy at 3700-2800 cm^-1^ band. Simple ds3T3A, ds3A3T, ds3C3G, ds3G3C, and complex ds3C3A, ds3A3C, dsCACGTG (canonical E-box), dsTTAGGG (telomere repeat), dsCTCGAG (XhoI), dsCTTAAG (AfIII), dsTCTAGA (XbaI) dsGAGCTC (SacI), dsGAATTC (BamHI), dsAGATCT (BgIII) in 0.1M NaCl solution were prepared at 100 pmole/μL as previously described. Each sample was treated with decagonal CEDS using each target sequence for 30 minutes at room temperature, and immediately analyzed by FT-IR (Perkin Elmer, USA). The positive control was treated with CEDS using a non-specific sequence (12A*-CEDS).

To exclude the O-H stretching of water, FT-IR analysis was performed together with distilled water (DW), 0.1M NaCl, and 1M NaCl solution as controls. Oligo-dsDNAs dissolved in 0.1M NaCl solution showed a slightly higher band than DW, but a lower band than 0.1M NaCl solution. However, oligo-dsDNA solution treated with CEDS showed significant increase in the IR absorbance band, while oligo-dsDNA solutions treated with nonspecific sequence 12A*-CEDS exhibited only slight increase compared to untreated control. When comparing Pyu and Puy oligo-dsDNAs treated with CEDS solutions, both Pyu and Puy dsDNAs showed increased IR absorbance at 3700-2800 cm^-1^ band compared to the positive control, and then Pyu dsDNAs tended to have more IR absorbance than Puy dsDNAs (Fig. 10).

**Fig. 10.**
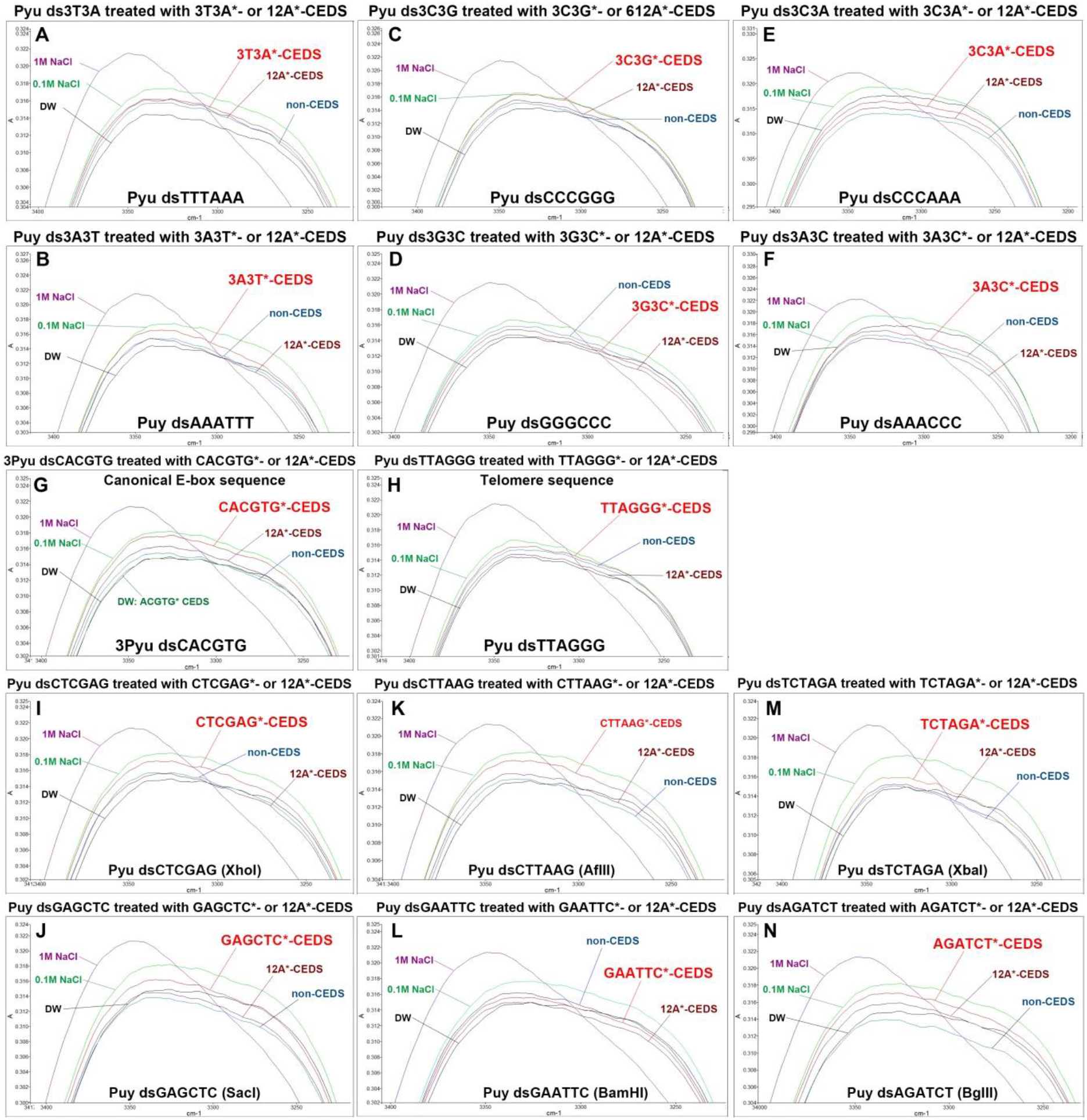
FT-IR analysis for CEDS effect on six bps simple and complex dsDNAs in 0.1M NaCl solution, comparing between each pair of Pyu and Puy oligo-dsDNAs, in comparison with the positive control (12A*-CEDS) and negative controls (no CEDS; DW, 0.1M and 1M NaCl solution).

## Discussion

This study is to develop the HBMR-based gene regulation system and propose decagonal CEDS with *in vitro* experiments using short (6-12 bps) oligo-dsDNAs. In particular, to avoid potential biohazards of alternating electromagnetic filed (*7, 8*), CEDS used a pulsed unipolar magnetic field with a frequency of 100 Hertz, strength of less than 30 Gauss, and duration of less than 30 min.

It was primarily found that CEDS can increase hybridization potential and induce unique conformation of target oligo-dsDNAs depending on CEDS time, which is reversible during resting time. Pyu oligo-dsDNAs are more responsive to CEDS than Puy oligo-dsDNAs. The CEDS increased the IR absorbance of oligo-dsDNA at 3700-2800 cm^-1^. Further investigation should be conducted to determine whether CEDS can affect the function of long (24 bps) oligo-dsDNA and plasmid DNA in the following study.

## Acknowledgments

We would like to express our gratitude to the late Professor Je Geun Chi and the late Dr. Soo Il Chung, who contributed to this research in part.

## References

1. S. K. Lee, D. G. Lee, Y. S. Kim, Development of Hydrogen Bonding Magnetic Reaction-based Gene Regulation through Cyclic Electromagnetic DNA Simulation in Double-Stranded DNA. arXiv:1210.7091, (2024); published online EpubJun 29 (10.48550/arXiv.1210.7091).

2. J. I. Jacobson, A look at the possible mechanism and potential of magneto therapy. Journal of theoretical biology 149, 97–119 (1991); published online EpubMar 7 (10.1016/s0022-5193(05)80074-8).

3. Q. Ren, J. Lu, H. H. Tan, S. Wu, L. Sun, W. Zhou, W. Xie, Z. Sun, Y. Zhu, C. Jagadish, S. C. Shen, Z. Chen, Spin-resolved Purcell effect in a quantum dot microcavity system. Nano letters 12, 3455–3459 (2012); published online EpubJul 11 (10.1021/nl3008083).

4. S. Sottini, E. J. Groenen, A comment on the pseudo-nuclear Zeeman effect. Journal of magnetic resonance 218, 11–15 (2012); published online EpubMay (10.1016/j.jmr.2012.03.009).

5. S. K. Kim, D. S. Lee, D. G. Lee, S. K. Lee, S. J. Choi, I. S. Song, Y. J. Lee, Y. W. Lee, S. C. Park, J. G. Chi, Genetic code is composed of pyrimidine-purine DNA segments as a basic unit displaying DNA base pair polarity. Medical Hypothesis and Research 5, 1–18 (2009).

6. S. K. Lee, Y. S. Kim, Y. W. Lee, J. G. Chi, S. C. Park, D. S. Lee, D. G. Lee, Method for visualization and polarity analysis of nucleotide using nucleotide base pair polarity, and program for genetic code analysis comprising thereof. Korean patent, KR101287040B101287041 (2013).

7. G. H. Kang, C. H. Lee, J. W. Seo, R. H. Sung, Y. H. Chung, S. K. Lee, Y. H. Suh, J. G. Chi, In-vivo study on the harmful effect of the extremely low frequency unipolar pulsating magnetic field in mice. Journal of Korean medical science 12, 128–134 (1997); published online EpubApr (10.3346/jkms.1997.12.2.128).

8. Y. S. Kim, S. K. Lee, Properties of Extremely Low Frequency Electromagnetic Field and their Effects on Mouse Testicular Germ Cells. International Journal of Oral Biology 35, 137–144 (2010); published online EpubSept 1 (https://db.koreascholar.com/Article/Detail/565).

